# Visual epidural field potentials possess high functional specificity in single trials

**DOI:** 10.1101/646612

**Authors:** Benjamin Fischer, Andreas Schander, Andreas K. Kreiter, Walter Lang, Detlef Wegener

**Affiliations:** Brain Research Institute, Center for Cognitive Science, University of Bremen, P.O. Box 33 04 40, 28334 Bremen, Germany; Institute for Microsensors, -Actuators, and –Systems, University of Bremen, P.O. Box 33 04 40, 28334 Bremen, Germany

## Abstract

Recordings of epidural field potentials (EFPs) allow to acquire neuronal activity over a large region of cortical tissue with minimal invasiveness. Because electrodes are placed on top of the dura and do not enter the neuronal tissue, EFPs offer intriguing options for both clinical and basic science research. On the other hand, EFPs represent the integrated activity of larger neuronal populations, possess a higher trial-by-trial variability, and a reduced signal-to-noise ratio due the additional barrier of the dura. It is thus unclear whether and to what extent EFPs have sufficient spatial selectivity to allow for conclusions about the underlying functional cortical architecture, and whether single EFP trials provide enough information on the short time scales relevant for many clinical and basic neuroscience purposes. We here use the high spatial resolution of primary visual cortex to address these issues and investigate the extent to which very short EFP traces allow reliable decoding of spatial information. We briefly presented different visual objects at one out of nine closely adjacent locations and recorded neuronal activity with a high-density, epidural multi-electrode array in three macaque monkeys. Using receiver-operating characteristics to identify most-informative data, machine-learning algorithms provided close-to-perfect classification rates for all 27 stimulus conditions. A binary classifier applying a simple max function on ROC-selected data further showed that single trials might be classified with 100% performance even without advanced offline classifiers. Thus, although highly variable, EFPs constitute an extremely valuable source of information and offer new perspectives for minimally invasive recording of large-scale networks.

## Introduction

The acquisition of spatiotemporally highly resolved activity from large neuronal populations has opened new possibilities for studying the distributed neuronal processes underlying the brain’s cognitive and executive functions (Varela et al., 2001; Siegel et al., 2012). A closer understanding of their spatiotemporal dynamics and discrimination of critical patterns within this activity offers promising perspectives for the diagnosis, treatment, and therapy of nervous system diseases (Engel et al., 2005; Murphy et al., 2016; Lebedev and Nicolelis, 2017; Parvizi and Kastner, 2018). Intracerebral recordings, using microelectrode arrays (Vetter et al., 2004; Suner et al., 2005) or microwire bundles (Williams et al., 1999; Nicolelis et al., 2003; Schwarz et al., 2014), provide access to single cell activity within a designated cortical region. Electrocorticography (ECoG), on the other hand, using subdural electrode grids, allows covering a large extent of the cortical surface and provides access to measurements at a mesoscopic scale.

ECoG was introduced during the early 1950s to identify epileptogenic zones prior to surgical intervention (Penfield and Jaspers, 1954), and later on became the gold standard for perioperative functional mapping (Palmini, 2006; Yang et al., 2014). ECoG arrays for clinical purposes mostly use electrodes with large diameter and large inter-electrode distances (Lesser et al., 2010; Schalk and Leuthardt, 2011). More recently, however, several research groups developed high-density arrays that allow analysis of neural signals on a finer spatial scale (Rubehn et al., 2009; Fukushima et al., 2014; Tolstosheeva et al., 2015; Schander et al., 2019). Because ECoG provides better spatial resolution, better signal-to-noise ratio, and higher frequency components (> 40 Hz) than electroencephalography (Ray et al., 2008; Ball et al., 2009), and constitutes a probably more durable recording approach than intracerebral electrodes (Chao et al., 2010), it has now become an increasingly interesting technique for both basic research on the dynamics of large-scale networks (Lewis et al., 2015) and brain-computer interfacing (Schwartz et al., 2006).

A major challenge in ECoG research is to extract useful information from short data fragments (i.e. single trials), due to the strong trial-by-trial fluctuations in both distributed and local brain activity underlying different perceptual and cognitive states (Fox and Raichle, 2007; Garrett et al., 2013), and the short time scales on which neuronal signals need to be classified for clinically relevant brain-computer interfaces (BCI). To increase the signal-to-noise ratio, ECoG arrays are mostly placed below the dura mater, to avoid attenuation of the signal by the barrier of the dura. Opening the dura usually does not constitute a limiting factor for peri-operative ECoG, whereas it may create a significant drawback for long-term recordings regarding invasiveness and related clinical complications (Lesser and Arroyo, 2005). As an alternative, multielectrode arrays may be placed on top of the dura to record the epidural field potential (EFP), thus minimizing the risk of side effects for chronic applications.

In the motor domain, EFPs were successfully used to decode hand and arm movements (Flint et al., 2012; Shimoda et al., 2012; Marathe and Taylor, 2013) and accurate decoding of motor commands with epidural signals was suggested as an important next step for viable long-term BCIs (Flint et al., 2017). Yet, little is known about the selectivity and reliability of EFPs from other cortical domains and how specific single-trial EFPs reflect the local functional circuitry.

We here make use of the high spatial resolution of primary visual cortex (V1) to investigate the functional specificity of single-trial EFPs recorded with a high-density, epidural multielectrode array. Because visual stimulation elicits clearly localized neuronal activations, V1 is particularly suited to study whether these are preserved in the EFP response. We show that despite its obvious limitations, the EFP allows close-to-perfect classification of the spatial information of nine closely adjacent stimuli at the single-trial level. The results suggest a strong potential of EFPs for both the trial-wise investigation of neuronal processes during sensory and cognitive processing and the further development of minimally invasive clinical approaches.

## Material and Methods

### Subjects

All experimental and surgical procedures followed the *Directive 2010/63* issued by the European Commission and the *Regulation for the Welfare of Experimental Animals* issued by the Federal Government of Germany, and were approved by the local authorities. EFP recordings were performed in three male macaque monkeys *(Macaca mulatta),* 13, 11, and 11 years old. All monkeys were familiar with laboratory standard procedures and a dimming task at fixation, and took part in additional projects not reported here. Monkey 1 (M1) and Monkey 3 (M3) were separately housed in indoor compartments, in visual and auditory contact with other animals. Monkey 2 (M2) was housed in a group of four animals, in a large indoor room with daily access to an equally large outer compartment. All compartments were enriched by a manifold of monkey toys, puzzles, and climbing opportunities. During training and recording sessions, monkeys received water or diluted fruit juice as reinforcer for performing the behavioral task in the recording chamber. At non-recording days, they received free fruits and liquid in their home compartment. Health and well-being was checked by daily monitoring and regular visits by veterinarians, and body weight was checked several times the week.

### Surgical procedures

For anchoring a headholder and the connector of the electrode array, all monkeys were implanted with a cap of acrylic cement (Palamed/Paladur), fixed by cortical screws to the animals’ skull, following a previously published protocol (Wegener et al., 2004). The EFP array was implanted after monkeys had been familiarized with the task and stimulus conditions reported below. Surgeries were performed under strictly sterile conditions. Anesthesia was initialized by a mixture of Ketamine/Medetomidine and was maintained by Propofol and Remifentanil, and supplemented by Isoflurane. Post-operative analgesia was maintained by Carprofen. For epidural implantation of the multi-electrode array over V1, the stereotaxic location of the lunate sulcus was estimated prior to surgery based on structural MRI scans. A trepanation over the left hemisphere was made using an ultrasonic bone cutting device with a 0.3 mm cutter, ranging from a few mm anterior the lunate sulcus to the occipital ridge, and from 2 mm lateral the midline to the lateral surface of the skull. The trepanation was extended in frontal direction into a 10 * 15 mm^2^ stripe to place the flat ribbon cables connecting the electrodes with OMNETICS connectors (MSA-Components, Attendorn, Germany). After the array was placed in its desired position, the location of some spatially characteristic electrodes was stereotactically determined and/or photographed. Ribbon cables left the cranium through a small gap in the skull. The trepanation was closed by reinserting the bone piece (kept in Ringer lactate during the meantime) and kitting with bone cement, and additionally fixing the bone flap with a stripe of titanium, anchored by cortical screws. OMNETICS connectors were placed in a trough of the Paladur cap, filled with two-component glue (UHU 300 endfest). Finally, a frame to hold a protective cap covering the connector was anchored. Additional ground electrodes were implanted over frontal cortex.

### Data Acquisition

The multielectrode array (Fig. 1*A*) was developed, designed, and manufactured at the University of Bremen (Schander et al., 2019). The design purpose was to match the size and shape of the macaque brain’s occipital surface for covering V1/2 and part of V4 in one hemisphere with a high number of electrodes. Each array consisted of 202 gold electrodes with 560 μm diameter, arranged in a hexagonal pattern with 1.8 mm center-to-center distance. The reference electrode was placed on top of the array and faced the skull. Electrodes were coated with poly(3,4-ethylenedioxythiophene) polystyrene sulfonate (PEDOT:PSS) to reduce the electrode/tissue impedance (Pranti et al., 2018). The substrate material was the polymer polyimide with a total layer thickness of 10μm. Ground electrodes over frontal cortex were coated with PEDOT: PSS, too.

**Fig. 1.**
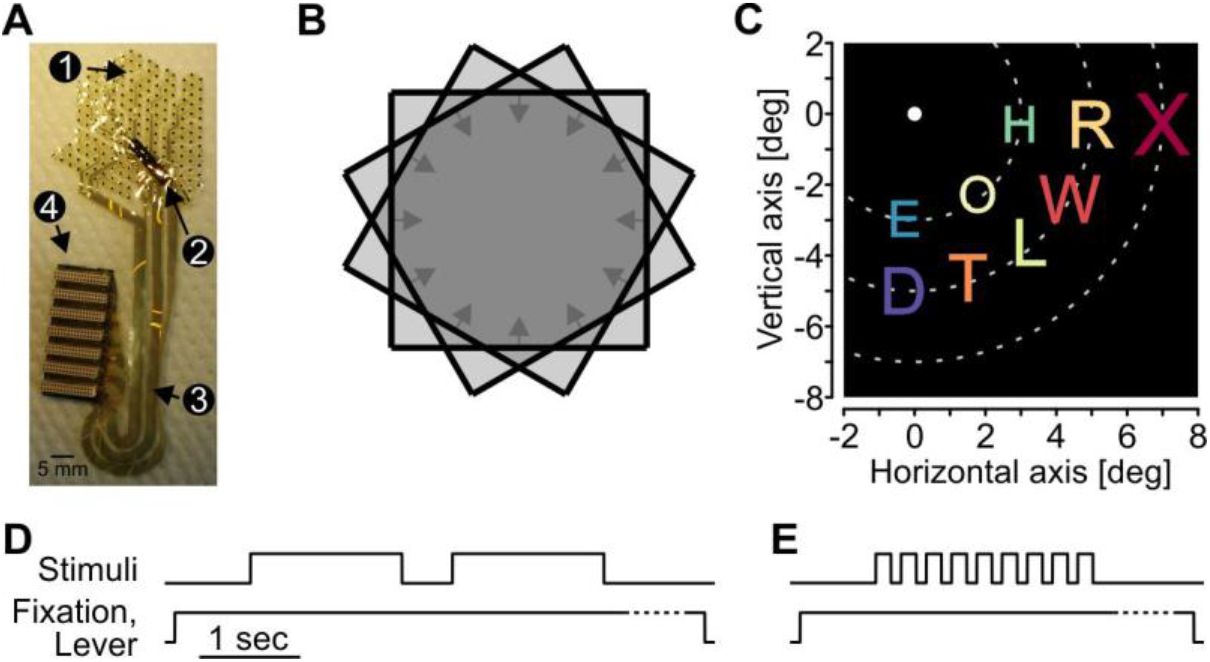
Multielectrode ECoG array and visual stimulation. A: Multielectrode array with 202 PEDOT:PSS-covered gold electrodes (1), reference electrode (2), conductor tracks (3), and OMNETICS connectors (4). B: Scheme of bar trajectories for the ERF mapping procedure. Black lines, differently oriented bar stimuli; arrows, motion directions; light grey-shaded areas, areas covered by motion trajectories; dark grey-shaded area, area of overlap of all trajectories. *C*: Spatial stimulus arrangement in the letter presentation procedure. Dotted lines indicate iso-eccentricity lines used for stimulus placing and were not shown on the display. *D - E:* Time course of trials in the mapping (*D*) and letter presentation procedure (*E*). During the mapping procedure either one (M1, M3) or two bars per trial (M2, as shown in sketch) were presented. Dotted lines indicate randomized period at end of trial, after disappearance of visual stimuli.

Recordings were performed with a sampling rate of 25 kHz. Pre- and main amplification (x10, x5000) and filtering (1 Hz – 5 kHz) was performed using equipment from Multichannel Systems (Reutlingen, Germany). The EFP signal, the analog eye position channels from a custom-made video-oculography system, and a 50 Hz power line-derived signal were stored on a computer for offline analysis. Data were recorded in multiple sessions within up to several weeks. The first session was recorded two (M3), five (M1), and eighteen (M2) weeks after array implantation.

### Visual stimulation and behavioral task

Visual stimuli were presented on a 21 inch cathode ray tube monitor (resolution: 1152 * 864 px, refresh rate: 100 Hz), placed 70 cm in front of the animal. Stimulation consisted of an automatic mapping procedure to characterize the size and location of EFP receptive fields (ERFs), and rapid presentation of nine different letter stimuli in the lower right quadrant of the visual field to investigate the spatial selectivity of single-trial EFP traces. The ERF mapping procedure was similar to a method recently introduced by Fiorani et al. (2014), described in detail in Drebitz et al. (2019). Briefly, an array of twelve oriented moving bars (size: M1: 0.24 x 16.1 deg; M2 and M3: 0.24 x 24.2 deg) was placed over the lower right quadrant (centered at 8.1/-4.0 deg for M1, and 2.4/-3.6 deg for M2 and M3). Bars were shown against a gray or black background (luminance: 2.5, 0, 0 cd/m^2^), with suprathreshold contrast (4.15, 7.85, 7.85 cd/m^2^). Each bar moved along a trajectory of fixed length (20.2 deg, 19.4 deg, 19.4 deg) in one out of twelve directions, spaced by 30 deg, with a velocity of 8.1, 12.9 and 6.5 deg/sec for M1, M2, and M3, respectively. The bars’ motion trajectories formed a regular, twelve-sided area of overlap with polygonal shape (Fig. 1*B*).

To investigate spatial stimulus selectivity of single-trial EFPs, differently colored, isoluminant (10 cd/m^2^) stimuli were shown at one out of nine adjacent locations in the visual quadrant covered by the electrode array. Background luminance was the same as during the RF mapping procedure. Stimuli consisted of the letters D, E, H, L, O, R, T, W, and X in Arial font. Letters were placed on three imaginary rings around the center of the screen (Fig. 1*C*), with a center distance of 3.5, 5.9, and 8.2 for M1 and 3, 5, and 7 deg for M2 and M3. Letter stimuli in the inner imaginary ring had a height of 1.25 deg for M1, and 0.95 deg for M2 and M3. Stimulus size on the middle and outer imaginary ring increased by factors of 1.2 and 1.5, to account for the increase in receptive field size with increasing eccentricity (Hubel and Wiesel, 1974). During presentation of visual stimuli, monkeys performed a dimming task at fixation. Monkeys were required to initiate a stimulus sequence by gazing at a central fixation spot (0.2 * 0.2 deg) and pressing a lever, and to respond to the dimming of the fixation spot by releasing the lever within 200 to 750 ms. Lever release outside the response window and eye movements of more than 1 deg away from the fixation spot caused termination of the trial. The temporal sequence in the mapping and letter presentation procedure is depicted in Figures 1*D* and *E*, respectively. Each stimulus sequence started with a blank screen of 750 ms duration, to provide a baseline period without visual stimulation for baseline correction of responses. During the mapping procedure, either one (M1, M3) or two bars after each other (M2) were shown, depending on the length of the motion trajectory. If two bars were shown in a single trial, they were separated by a blank interval of 500 ms. Motion directions were chosen randomly and each direction was presented at least ten times. During letter presentation, all nine stimuli were shown per sequence, each for 150 ms, separated by a blank interval of 100 ms, in pseudo-random order. For both the mapping procedure and the letter presentation, dimming of the fixation point occurred during an interval 0.1 to 1.1 sec after disappearance of the last object shown. Subsequent sequences were separated by an intertrial interval of maximally 1.5 sec. In M1 and M3, FP dimming occurred during stimulus presentation in about 5% of the trials (catch trials). These trials did not enter data analysis.

### Data Analysis

#### Preprocessing

All recorded signals were low-pass filtered (FIR filter in forward and backward direction, 150 Hz cutoff frequency) and down-sampled to 1 kHz. If applicable, 50 Hz noise was removed based on the phase of the recorded power line signal. Electrodes located anterior to the lunate sulcus and electrodes with no or strongly noise-disturbed signals were excluded. The final database included 88 (M1), 174 (M2), and 137 (M3) electrodes delivering a significant ERF (see below) and covering V1.

#### EFP signals

We use the term EFP to denote single-trial signals at any given electrode following the onset of a stimulus. At each electrode, EFPs were baseline-corrected by subtracting the pre-stimulus signal amplitude (averaged over time and trials) from each trial’s stimulus response. EFPs were then smoothed with a Loess filter with λ = 2 and α = 0.04. All response features of the EFP that were used for classification (see below) were computed either from the whole time course of the trial between 25 and 175 ms after stimulus onset, or alternatively, from a shorter time period estimated by ROC analysis (see below). EFP features that were tested were e.g. maximal amplitude and maximal absolute amplitude, averaged over the respective time bins, and negativity (most negative peak during the respective time period). Alternatively, the time course of the EFP modulation during the entire trial was used as feature.

#### Wavelet transformation

To obtain spectral information, EFPs of each electrode and stimulus sequence were wavelet-transformed using Morlet wavelets (Torrence and Compo, 1998), defined as:

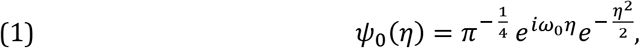

where *ω*_0_ refers to the non-dimensional frequency (set to six to cover 5-160 Hz in 35 exponentially increasing steps) and *η* refers to the non-dimensional time parameter. The wavelet transform is defined as:

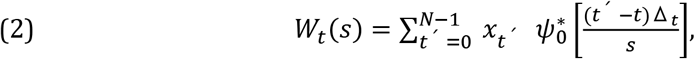

where *x_t_* indicates the time series, Δ_*t*_ the time step, *s* the scale of the wavelet, and * denotes complex conjugation. Wavelet transformation was performed on raw EFPs, as obtained after preprocessing. The resultant complex numbers *W_t_* can be written as:

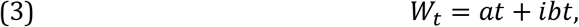

where *a* is referring to the real part *Re*{*W_t_*} and *b* to the imaginary part *Im*[*W_t_*], both of which were used as candidate features for classification (see below). Wavelet power *WP_t_* for each time bin *t* was calculated by:

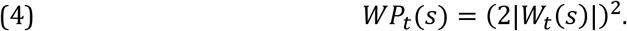

For each frequency band, baseline correction was done by subtracting mean baseline *WP* from the mean *WP_t_* during stimulus presentation, and subsequently dividing through it. Time-trigger information was used to cut the continuous data into stimulus-specific data fragments.

Features extracted from the time course of the wavelet-transformed data were e.g. mean power in a given time-frequency range, their real and imaginary part, and phase angle. Mean power was calculated by averaging over all bins of a given frequency range of the entire trial, from 25 to 175 ms post-onset, or alternatively, over a limited time-frequency window estimated by ROC analysis (see below). Mean real or imaginary parts of *W_t_* were computed accordingly. The phase angle of a given time-frequency range was expressed as the circular mean over the phase angles of individual bins, using the ‘CircStat’ toolbox for Matlab (Berens, 2009). Alternatively, the trial’s time course of the mean power between 60 and 150 Hz (broadband gamma power, BGP) was used as feature for classification (see below).

#### Receptive field mapping

Prior to computation of ERF maps, data with extraordinarily high continuous activity throughout the trial was excluded using an automated trial-rejection procedure. Subsequently, ERF calculation for each electrode was performed by first averaging mean *WP_t_* across the BGP frequency range of all trials of a given motion direction. The response *R* at each location (*x, y*) was expressed as the geometric mean over the Z-transformed responses to the *n* = 12 motion directions at the time the bars crossed that location plus a constant value of *c* = 80 ms to account for response latency. ERFs were defined as area of *R_map_* with Z-scores > 1 and minimal size > 1 deg^2^ (Fig. 2*A, B*). If more than one area was found, only the largest was chosen. A detailed description of the procedure is provided in Drebitz et al. (2019). As a measure for the strength of the response relative to the background activity we calculated the signal-to-background ratio (SBR) of ERFs by weighting power inside the ERF against power of the background:

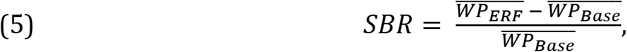

where 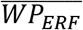 and 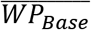 indicate mean power inside and outside ERF, respectively. All analytical procedures taking into account the ERF center refer to the center of activation (peak activation).

**Fig. 2.**
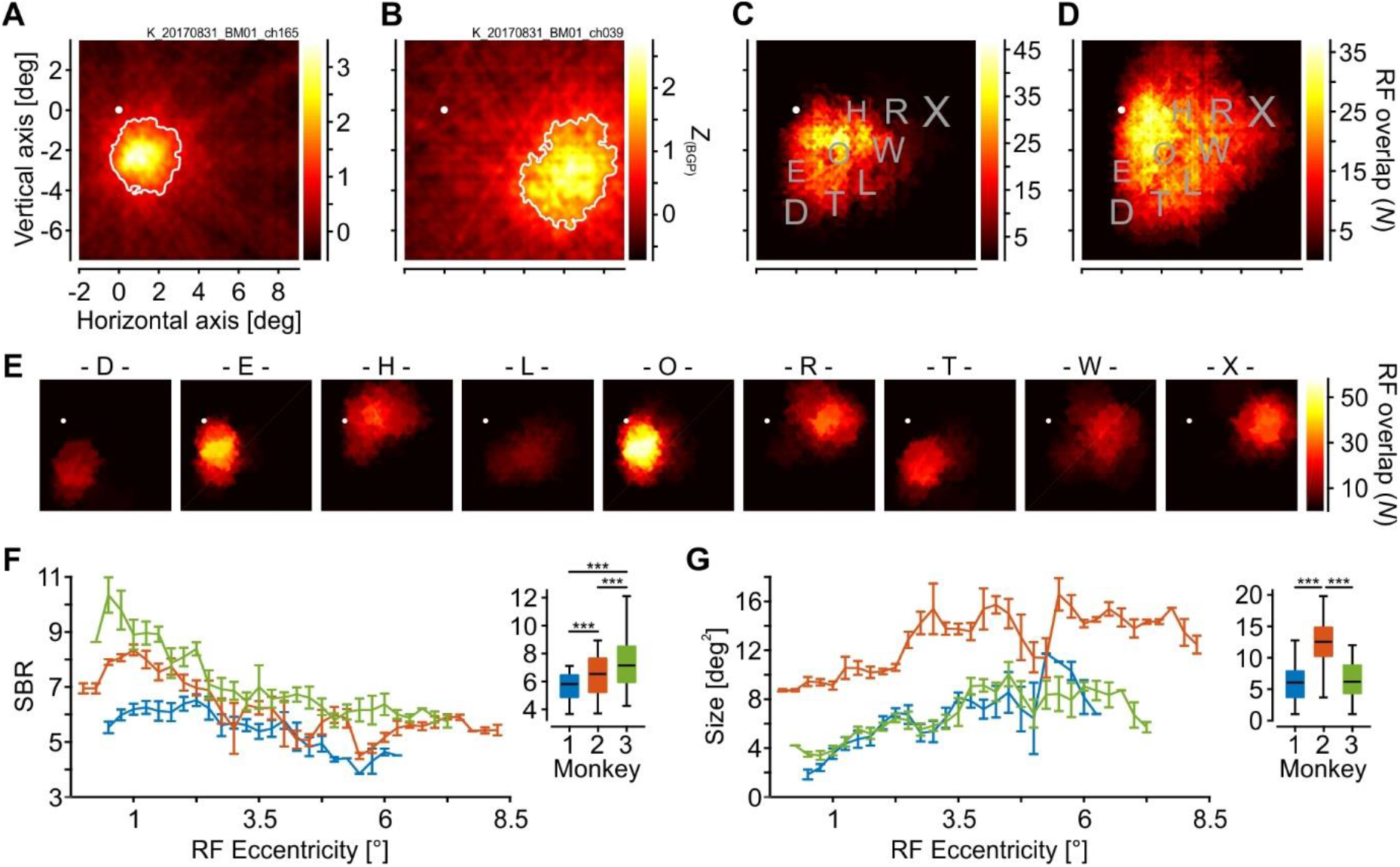
ERF coverage of visual fields and basic ERF characteristics. *A*: Visual response map of one example electrode in M2 showing well-defined area with significant evoked response (ERF, white border), obtained from an automated bar-mapping procedure. *B*: Single, rather large ERF mapped from another electrode in M2. *C - D:* Map of merged ERFs from M1 (*C*) and M3 (*D*). Color code indicates the number of overlapping ERFs at each position. Letter stimuli are shown for reference only and were not used for mapping ERFs. *E*: Same as in *C* and *D*, but referenced to the location of individual letter stimuli, as used in subsequent procedures. For each letter position, all ERFs with center coordinates within 2 deg from the stimulus center were considered. Color axis is identical for all maps. *F - G:* Signal-to-background ratio (*F*) and size (*G*) as function of ERF center eccentricity (line plots) and averaged over all electrodes per animal (box-plots). Horizontal black lines in box-plots indicate median, boxes indicate the 25^th^ and 75^th^ percentile, and whiskers show full range of data. Error bars in line plots indicate S.E.M. White spots in *A - E* indicate fixation point. Scaling of x- and y-axes in *B - E* is indicated in *A*. Stars in *F* and *G* indicate significant differences with *P* < 10^−3^.

#### Support vector machines

Investigation of the spatial selectivity of single-trials was performed by training support vector machines (SVMs) on data from the letter presentation procedure. No trial rejection for exclusion of data with artifacts was performed, to more closely simulate clinical conditions. Throughout the remainder of the paper, we use the term ‘trial’ to denote the response to a single stimulus in a stimulus sequence, between 25 and 175 ms after stimulus onset. Multiclass classification was based on radial basis function kernel, one-vs.-rest SVMs, as implemented in “libsvm” (Chang and Lin, 2011). For classifying single-trials, we randomly chose 50 trials per stimulus condition for training one SVM for each of the nine stimulus conditions. If not stated differently, SVM parameters c and γ were chosen optimal for each training set using grid search with cross-validation (c represents the cost function and can be thought of as the penalty for misclassification during training, γ is the free parameter of the Gaussian radial basis function and can be thought of as the spread of the decision region). For each of the nine stimulus conditions, one test trial, not used in any of the training sets, was assigned to the stimulus with the highest probability for class identity. This procedure was repeated for each of the trials in the database. Per monkey, stimulus, and (where applicable) electrode, SVM classification performance was calculated by comparing assigned vs. real identity over all test trials.

#### Single-trial classification based on time course

To investigate the stimulus specificity of single trials, we first tested a signal’s time course at individual electrodes of the three arrays. This was done by training SVMs as described above, using first, the baseline-corrected EFP time course and second, the BGP time course of the wavelet-transformed data, recorded between 25 and 175 ms after stimulus onset. To keep computations within reasonable time, we initially used a limited range for SVM parameters c and γ for single-electrode classification. Based on these results, we then selected the best-performing electrode per stimulus (Best1 condition), or the concatenated response of the best three or best five electrodes per stimulus (Best3 (5) condition) for optimized classification, using a wider range of parameters c and γ (as explained earlier). We additionally concatenated the response vectors of all the best-performing electrodes into one long vector (Best1paired condition).

#### Single-trial classification based on distributed activity

As an alternative for using the time course of the signal, each trial was expressed as the distributed activity over the entire array. To this end, each electrode’s time course was collapsed to a single value (using different response features, as e.g. mean EFP amplitude or mean BGP power over time), such that each trial was now represented by a vector of length *N*, where *N* corresponds to the number of electrodes (Fig. 4*B*). Training and testing of SVMs was performed as described earlier.

#### Initial selection of informative response features

T-distributed Stochastic Neighborhood Embedding (t-SNE) (van der Maaten and Hinton, 2008) was used for initial visual exploration of various candidate features (e.g. power in a specific frequency range). T-SNE is an unsupervised, non-linear machine learning algorithm for dimensionality reduction, improving clustering of high-dimensional data in low-dimensional space by minimizing the distance of neighboring data points and maximizing the distance of distant points. The Perplexity parameter was set to 30 and maximal number of iterations was limited to 1000.

#### Selection of most-informative time (-frequency) windows

Much of the variability of neuronal responses is explained by internally-driven activity (Fiser et al., 2004), and intrinsic signal fluctuations may cause stronger response modulations than does the visual input (Lee et al., 2019). To isolate informative, stimulus-specific time periods during stimulus responses from nonspecific intrinsic fluctuations in the signal, we applied Receiver-Operating Characteristic (ROC) analysis. For each monkey, ROC curves were computed by testing the trials of one condition against the trials of all other conditions for each individual time bin. To improve reliability, temporal resolution of the data was averaged within successive bins of 5 ms length. The time course of the variance of the area-under-the-ROC curve (AUC) was then used to select putatively most-informative time bins. For individual time bins, AUC variance is low in the absence of a visual response to any of the stimuli, when the ROC classifier is guessing and AUC values for all nine stimuli are close to 0.5. In contrast, AUC variance is increasing in the presence of a selective visual response, when there is good classification for one (or a small number) out of all stimulus conditions. Empirically, we found AUC variance to be more selective and more indicative of putatively informative time bins then were other measures like mean or maximal AUC.

The time-resolved ROC analysis was performed separately for the EFP and for the power, phase angle, *Re*{*W_t_*}, and *Im*{*W_t_*} of the individual frequency bands of the wavelet-transformed data. Subsequently, for each of the different measures, the grand variance of AUC values was calculated bin by bin over all stimulus conditions and electrodes of each array, such that for each of the measures high variance indicates bins with strong signal differences between electrodes and conditions, i.e. presumably more distinct responses to the test stimulus. All values were then *Z*-transformed to allow for a single selection criterion across features and electrode arrays. Aggregates of three or more bins with a Z-score > 1.5 were chosen for further analysis. If more than one such aggregate was found for a given feature, the one with the highest peak variance was selected. Per electrode, final features were expressed as means over the selected time (or time-frequency) bins (e.g. mean power in a specific time-frequency cluster).

#### Selection of most-informative electrodes

In analogy to identifying the most-informative time bins, we identified electrodes carrying most of the information in the distributed activity vectors using the same ROC-AUCs. For each electrode, AUC values of the most-informative time (or time-frequency) bins were averaged for each stimulus and expressed as absolute deviation *Δ* from 0.5 (guessing). Electrodes were then ranked from max to min *Δ* for each stimulus, resulting in nine different, stimulus-specific rank orders. Most-informative electrodes were selected based on their ranks for the response feature under investigation. Alternatively, to test the combination of different features, ranking could be based on the mean AUCs over these features.

### Experimental design and statistical analysis

For each of the three monkeys, all electrodes delivering a significant visual response during the ERF mapping procedure were considered. Statistical analysis was performed by first testing for normal distribution of the data using Shapiro-Wilk tests. Non-parametric tests were used if the Shapiro-Wilk test rejected the null hypothesis of normal distribution at the α = 5% significance level for at least one of the data sets. Effect size *ω*^2^ (Hays, 1963; Lakens, 2013) for both parametric and non-parametric ANOVAS (Kruskal-Wallis) was calculated by:

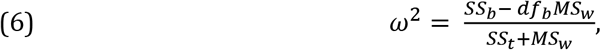

with all values for calculation derived from the statistics of the preceding test. *ω*^2^ ranges between 0 (no effect) and 1. For Wilcoxon signed rank tests, effect size *R* was calculated by:

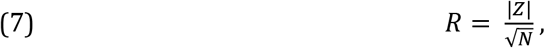

where *Z* was derived from the Wilcoxon test statistics and *N* indicates the total number of samples. As a benchmark, effect size is considered to be small for *R* = ~0.1 and large for *R* >= ~0.5 (Cohen, 1988; Field, 2013). Where applicable, post-hoc analyses were corrected for multiple comparison using Tukey’s honest significant difference criterion.

## Results

The goal of the study was to assess the spatial specificity of visually driven EFPs from primary visual cortex, and to quantify whether, and how reliable, spatial information can be drawn from single trials, lasting no longer than 150 ms. To this end, we recorded visual responses from three monkeys with epidurally implanted multielectrode arrays over V1 and visually stimulated at nine close-by locations in one visual quadrant. In the following sections, we will first analyze size and coverage of EFP-RFs (ERFs), and then investigate the classification rate at single electrodes and at groups of selective electrodes, using machine learning algorithms trained on the time course of the signal. We then continue by analyzing the distributed activity over the whole array and apply ROC analysis to identify most-informative response features, time and time-frequency windows, and electrodes. Based on the results of the ROC analysis, we finally isolate the most-informative data and show that these allow to draw spatial information from single EFP trials with extremely high reliability.

### ERF coverage of the epidural multielectrode array

To estimate the coverage of the visual field by the microelectrode array, we analyzed the location and size of areas with significant activation, using an automated bar-mapping procedure. The bar mapping procedure elicited strongest, and spatially most selective, responses in the broad gamma-frequency range (60 – 150 Hz, BGP). Accordingly, ERFs were defined by their mean activation in the BGP range (> 1 Z-score) and a minimum size (> 1 deg2). All analyses that follow were carried out on this set of electrodes with a significant ERF (88, 174, and 137 recording sites in M1, M2, and M3, respectively).

At some electrodes, responses to visual stimuli had a high signal-to-background ratio (SBR) and ERFs were of rather limited size (Fig. 2*A*). At others, ERFs were larger and frequently covered the location of several of the stimuli intended for later visual stimulation (Fig. 2*B*). Based on the size and location of ERFs, we constructed activation maps quantifying the number of ERFs covering each location within the visually stimulated field. These maps show dense sampling at some locations and sparse sampling at others (Fig. 2*C – D*, data taken from M1 and M3), likely due to cortical magnification at fovea, actual coverage of the cortical surface by the array, and regular arrangement of array electrodes. As a consequence, the representation of the nine letter stimuli is expected to vary significantly over the array, not only with respect to location but also regarding the number of electrodes carrying the signal and the separability of their representation from the representation of neighboring stimuli. This is illustrated in Figure 2*E*, showing putative ERF activation maps for individual letter stimuli (data taken from M2). Maps were built by considering all ERFs with a center within two degrees from the center of the respective letter stimulus. Based on the results of the bar mapping procedure, these maps predict that some stimuli will be represented by a large number of electrodes (e.g. letters E and O), while others will be represented by a few electrodes only (e.g. letters D and L), due to the size of ERFs or location of a stimulus at the edge of the visual field covered by the array. The results also predict that some stimuli are likely to activate electrodes with strongly overlapping ERFs, or activate the same electrodes (e.g. letters D and T). Thus, size and distribution of ERFs indicate that some letters are likely to yield better classification results than others. Additionally, we analyzed SBR and size of ERFs as a function of ERF eccentricity (using a sliding boxcar window of 2 deg width, shifted in steps of 0.25 deg). For each array, SBR was significantly larger at more foveal locations and decreased towards the periphery (least-square linear regression, slope = [−0.09 −0.08 −01.4], all *F* > 51.8, *P* < 10^−7^, adjusted *R*^2^ > 0.61) (Fig. 2*F*). Between arrays, SBR was significantly different (Kruskal-Wallis, *χ*^2^ = 55.41, *P* < 10^−12^, *df* = 2, *ω*^2^ = 0.135), with SBR being larger in M3 than in both M1 and M2 (both *P* < 10^−3^), and also being larger in M2 as compared to M1 (*P* = 10^−4^). For ERF size, we found a significant increase towards the periphery for M1 and M2 (slope = [0.29 0.17], *F* > 33.1, *P* < 10^−5^, adjusted *R*^2^ > 0.49), while the fit for M3 was too poor (adjusted *R*^2^ = 0.09) to draw a conclusion (Fig. 2*G*). Median ERF size over all electrodes of the arrays was 6.1, 12.5, and 6.2 deg^2^ in monkeys M1 to M3. Statistically, this was a significant difference between arrays (Kruskal-Wallis, *χ*^2^ = 217.45, *P* < 10^−47^, *df* = 2, *ω*^2^ = 0.543), and ERF size was larger at array electrodes in M2 than in M1 and M3 (both *P* < 10^−9^), while it was not different between M1 and M3 (*P* = 0.686). Thus, for the 3 * 9 stimulus conditions in the experiments that follow, analysis of ERF parameters suggests considerable differences for the representation of the letter stimuli across individual arrays as well as between monkeys. The variance in ERF parameters probably reflects differences in the distance between electrodes and cortical tissue, either due to technical reasons like imperfect fitting of the array to the dural surface, or due to variances in dura thickness or volume of subdural space. High variability in ERF size, SBR, and coverage is likely influencing the reliability by which single trial information can be read out from individual electrodes and (regarding the general usability of EFPs for basic and clinical science) addresses the question whether it is possible to find a robust approach to deal with this.

### Time course-based classification of single trials

To investigate stimulus specificity of singletrial signals recorded with epidural electrodes, we flashed visual letter stimuli at nine closely adjacent locations in the lower right quadrant of the visual field. Stimuli at each location were shown for 150 ms, separated by a blank screen of 100 ms length (Fig. 1*C* and *E*). They were isoluminant but differed in color and shape. For two example electrodes and three letters each, Figure 3*A* shows the EFP between 25 and 175 ms post-onset of ten randomly drawn trials per condition, and the corresponding ERF. Note that ERFs were recorded in a separate session. The electrode in the top panel (taken from M1) represents a class of recording sites with rather unspecific and strongly fluctuating responses, with no clear differences between responses to stimuli covering the ERF and stimuli outside the ERF. In contrast, the electrode in the lower panel (taken from M2) represents a class of recording sites with clearly modulated EFP traces in response to the stimulus covering the ERF. Yet, trials with a similar time course were acquired also in response to close-by stimuli. Figure 3*B* shows two more examples using wavelet-transformed derivates of the EFP. Time-frequency plots averaged over all trials per condition reveal that most of the signal’s modulation occurs at gamma-frequencies > 60 Hz, as covered by the BGP range used for ERF analysis. At some electrodes where the ERF covered a single stimulus, the BGP response was highly selective (Fig 3*B*, upper panel). At other electrodes, however, single trial responses were more variable in strength and timing, and very similar responses were elicited by neighboring stimuli and occasionally also by stimuli presented at clearly distant locations (Figure 3*B*, lower panel).

**Fig. 3.**
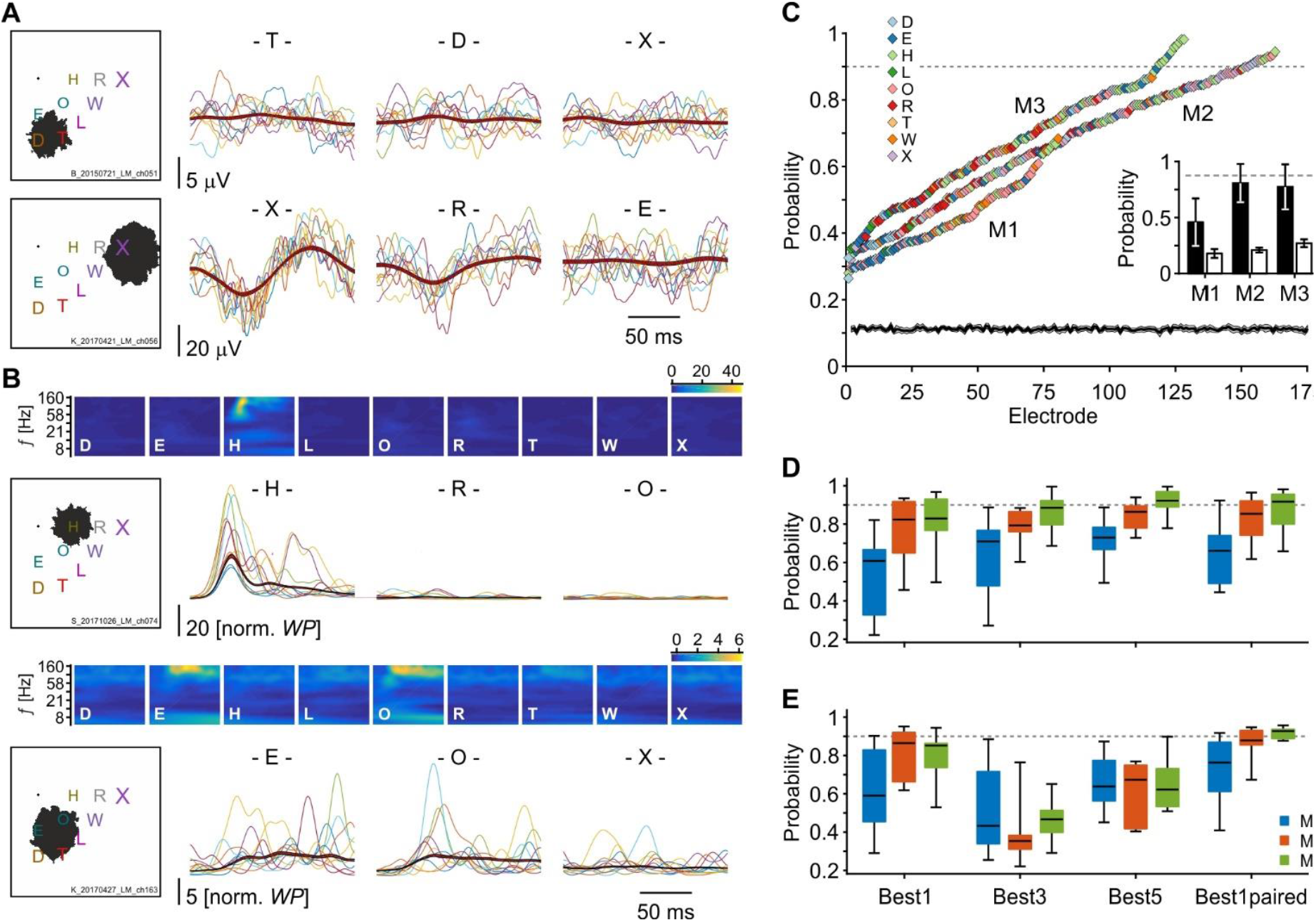
Single-trial classification performance at single electrodes. *A*: Stimulus location and ERF (black areas) for 2 example electrodes (left panels), taken from M1 (top) and M2 (bottom). For each electrode, panels to the right show the EFP signal of ten randomly drawn trials for three of the nine stimuli. Thick red lines indicate mean ± S.E.M over all trials of the respective condition. *B*: Same as in *A* but panels to the right show the BGP signal of ten randomly drawn trials for two more example electrodes, taken from M3 (top) and M2 (bottom). Top panels show the time-frequency plot of the same two electrodes in response to each of the nine stimuli, averaged over all trials, during the 25 – 175 ms time period after stimulus onset. Note the strong stimulus-induced modulation in the BGP range, and absence of modulation in other frequency ranges. Color code indicates normalized power. *C*: EFP-based single-trial SVM classification performance for all individual electrodes. Electrodes are sorted by probability for getting the true stimulus identity, individually for each animal. Colors indicate the identity of the stimulus that was most often correctly classified. Black bars in inlet indicate mean classification performance ± S.D. considering the best letter stimulus of each electrode, for each individual array. White bars indicate mean performance ± S.D. for all other stimuli at the same electrodes. Black solid line and grey shading at bottom of figure indicate mean chance level ± S.E.M., estimated by classifying M2 single trials with shuffled labels. *D* and *E*: Mean SVM classification performance for each of the three animals with optimized parameters c and γ, summarizing the results for both EFP (*D*) and BGP (*E*). SVMs were trained using either only the electrode with the highest performance for each of the nine letters (Best1), the concatenated responses of the best three or best five electrodes for each of the letters (Best3, Best5), or the concatenated responses of the nine best electrodes from the Best1 comparison (Best1paired). Box plot conventions as in Figure 2. Dashed lines in *C - E* indicate 90% performance level.

To investigate the information content of these signals, we trained support-vector machines (SVMs) on the time course from 25 to 175 ms of both the EFP and its wavelet-transformed BGP derivate, and tested SVM performance for each electrode separately. We classified a total of 6552 trials (2763, 1800, and 1989 trials from M1, M2, and M3, respectively). For the EFP (Fig. 3*C*), we found 13 (7.5%) and 11 (8%) electrodes in M2 and M3, respectively, performing at 90% or higher (see Material and Methods). Yet, these electrodes only represented four (M2) and two (M3) different stimuli, while maximal performance for four of the twelve remaining stimuli of these two monkeys and for all the stimuli in M1 stayed below 75%. Performance averaged over the best electrodes per letter stimulus was 46.1, 81.1, and 77.7% in M1 – M3, respectively. In comparison, mean performance of the same electrodes in response to all other letters was 17.9, 21.1, and 27.1% (Fig. 3*C*, inset). Using the BGP time course instead of the EFP trace improved the classification rates, such that now five (M2) and four (M3) stimuli were correctly classified at a performance level of 90% or higher. Yet, performance for half the stimuli of M2 and M3, and for all stimuli of M1 still remained below that level.

Because intracortical V1 signals are spatially highly specific, we considered a detection rate of 90% or better as the wished-for classification performance for EFPs. To achieve better classification performance, we therefore selected the best electrode for each of the nine stimuli and re-trained SVMs with optimized parameters c and γ. We also estimated classification rates using the best 3 and the best 5 electrodes of each stimulus, as well as the concatenated responses of the best single electrodes per stimulus (see Methods) (Fig. 3*D* and *E*). Statistical analysis revealed that the latter delivered the highest classification rate for both EFP (mean: 78.6%) and BGP (mean: 85.6%), and that BGP allowed for significantly better performance than the EFP (Wilcoxon signed rank, *Z* = 6.13, *P* < 10^−9^, *R* = 0.217, *N* = 399). Yet, 48% of the stimulus locations did not reach the 90% performance level, and 18.5% stayed below 75% correct. Thus, given the high variability of the signal, using the time course at single electrodes does not allow to sufficiently classify single trial information.

### Representation of single trials by patterns of distributed activity

As an alternative to single electrode-based classification, individual trials may be described as distinct patterns of distributed activity, due to the large cortical surface covered by the electrode arrays. The upper and lower panels in Figure 4*A* show the mean EFP and BGP amplitude, respectively, in response to stimuli at three different locations, averaged over about 100 trials of M1 (EFP) and M3 (BGP). It is obvious from these examples that the distinct, stimulus-specific spatial distribution of activity constitutes a valid source of information for classifying different stimulus conditions when considering averaged data. If similarly distinct patterns were present also in single trials, they may establish a more robust stimulus representation than the signals’ time course.

**Fig. 4.**
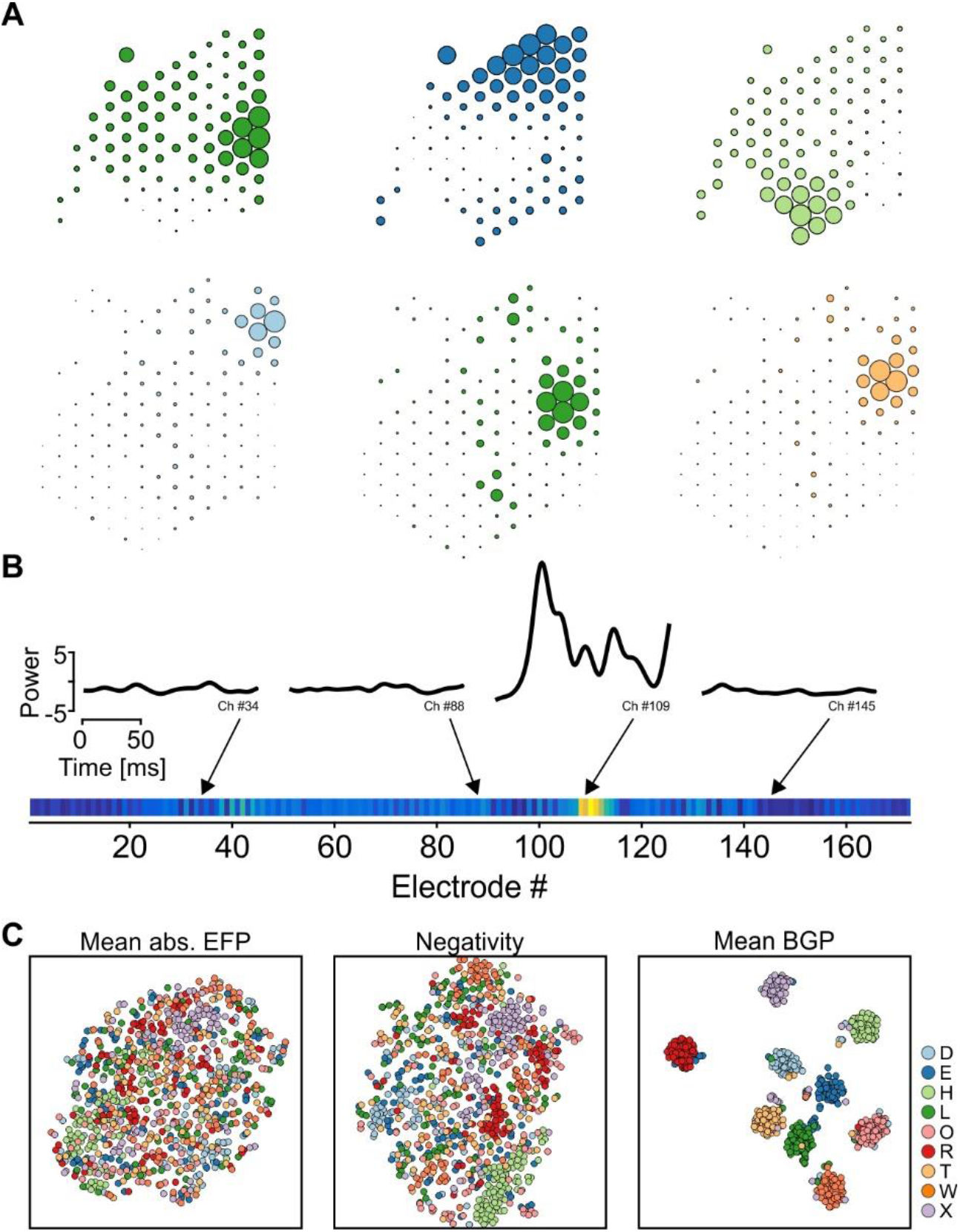
Spatial distribution of response features. *A*: Example plots of the spatial distribution of relative mean EFP over the array of M1 (top panels) and relative BGP over the array of M3 (bottom panels), averaged over the time period 25 to 175 ms post-onset of all trials of the respective stimulus. All values were squared for illustration purposes. Color code is indicated in *C*. *B*: Example for construction of trial vectors representing the distributed activity over the array (distributed activity vectors). For each electrode, the time course of the signal was collapsed to a single value (e.g. mean absolute EFP or mean BGP), or alternatively, only a single value was taken from the response (e.g. negativity). *C*: T-SNE plots for three different response features (data taken from M3), each represented as the distributed activity of a given stimulus feature over the entire array, as illustrated in *B*. Each dot represents a single trial. Well-defined clusters indicate good separability.

We therefore collapsed the signals’ time course at each electrode to a single value (e.g. mean EFP amplitude, negativity, mean power in a certain frequency range, etc.) and used the distribution of this feature over the entire array as the representation of a single trial (Fig. 4*B*). This way, each trial is described by a vector of length *N*, where *N* corresponds to the number of electrodes. We first visually explored different response features using t-SNE (van der Maaten and Hinton, 2008). T-SNE is a nonlinear, non-supervised machine learning algorithm for embedding high-dimensional data in low-dimensional space, providing a visual means to investigate whether data contain sufficient structure to distinguish different stimuli with some reliability. For the majority of features there was no or very weak clustering (Fig. 4*C*, left and middle panel), while for some we found distinct, well separated clusters (Fig. 4*C*, right panel). Thus, expressing a single trial as pattern of distribution of a given response feature over the array may allow to achieve higher classification reliability. At the same time, however, a significant number of trials was assigned to the wrong cluster even for overall well-clustering features, or clusters had considerable overlap. A possible reason for this is the inclusion of time periods containing random signal fluctuations rather than stimulus-specific information when considering the whole time course of the signal for feature extraction. Therefore, a more distinct representation of each stimulus might be achieved by getting rid of such non-informative trial periods.

### Identification of most-informative features and time bins across the entire array

To determine the stimulus selectivity of individual time bins, we performed a time-resolved ROC analysis to find the most-informative time periods to be used for stimulus classification. This was done separately for the EFP, the different frequency ranges of the wavelet-transformed data, phase angles, and both the real and imaginary part of the wavelet transformation, *Re*{*W*} and *Im*{*W*}. Prior to ROC, each signal’s time course was averaged into bins of 5 ms. Subsequently, for each feature, electrode, and time bin, we calculated the area-under-the-ROC curve (AUC). Figure 5*A* provides an example for the EFP at one example channel of M2, averaged over all trials per stimulus condition, and the ROC curves for three 5 ms-time bins, taken from the early, middle, and late response at that electrode. For the early and late time bins, the ROC curve for any of the nine stimuli was close to the diagonal, resulting in very similar AUC values around 0.5, and indicating poor separability of the stimuli. Yet, for the time bin around 55 ms the EFP signal showed a strong negative deflection in response to one of the nine stimuli, providing a rather large AUC value, while AUC values to all other stimuli remained around 0.5. The resultant distribution of AUC values possesses a higher variance than did the AUC distribution of the other two time bins (cf. inlets in Fig. 5*A*). We used this AUC variance to distinguish putatively informative from non-informative time bins, with high variance indicating stronger and potentially more stimulus-specific signal deflections than low variance. Following ROC analysis of all time bins and electrodes, we calculated the time-resolved grand average of AUC variance over all electrodes, and finally *Z*-transformed this to allow for better comparability across features and monkeys.

**Fig. 5.**
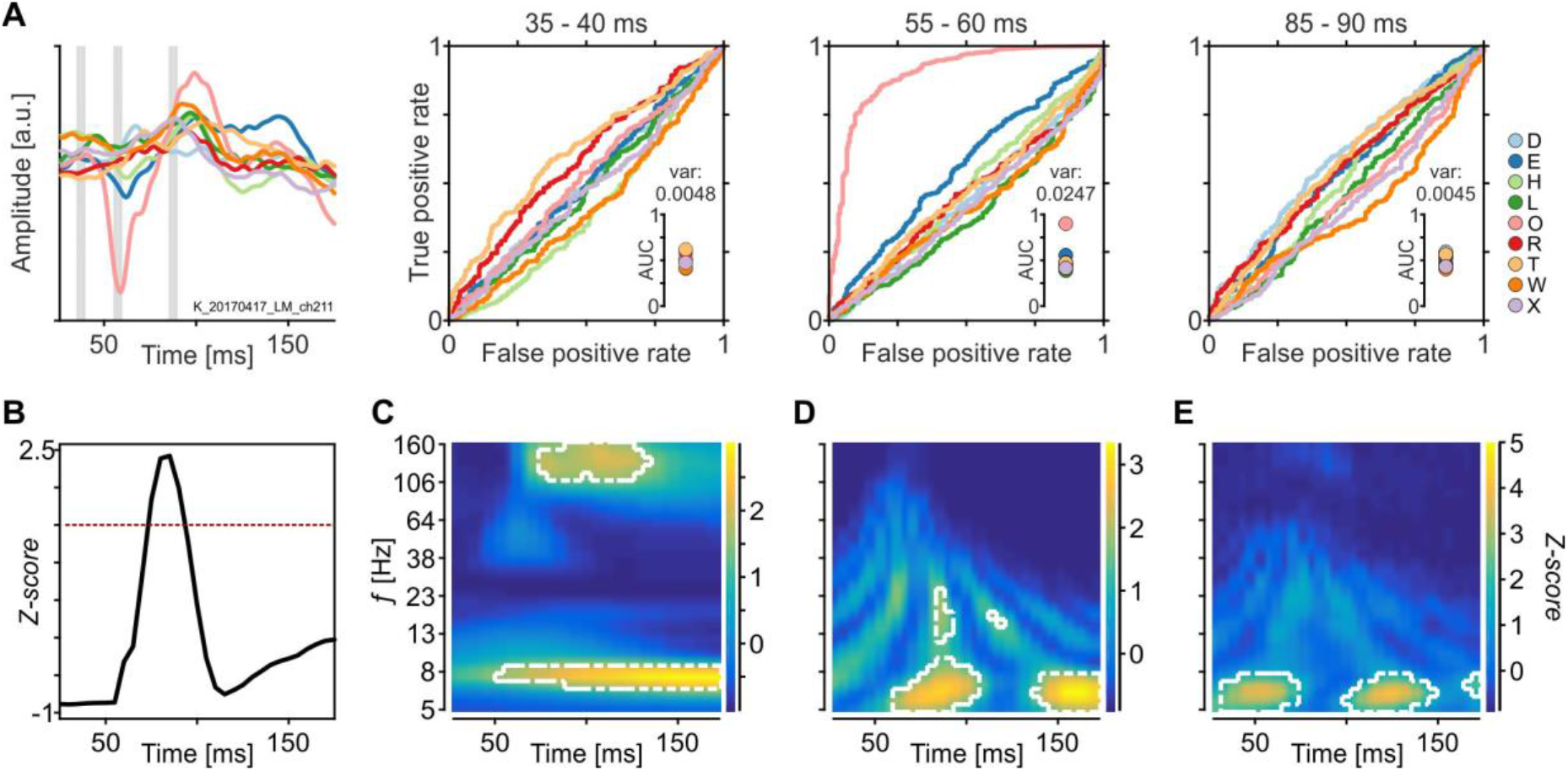
Identification of most-informative time epochs. *A*: Mean EFP traces to each of the nine stimuli at one example electrode from M2 (left panel), and ROC analysis of three 5 ms time bins as indicated by the grey-shaded areas in the left-most plot. Insets in ROC plots show distribution and variance of AUC values. *B: Z*-transformed grand mean of the variance over AUC values of M2, considering all electrodes, after bin-wise ROC analysis of EFP amplitudes. *C – E:* Same as in *B* but for time-resolved variance in power (*C*), real part of the wavelet coefficient (*D*), and phase angles (*E*) at 35 different frequency bands. Dashed lines in *B – E* indicate threshold at *Z* = 1.5.

For the EFP, the resultant time course of AUC variance shows a strong positive peak around 80 ms after stimulus onset. Applying a threshold of 1.5 labels a time window of about 25 ms as the putatively most informative epoch (Fig. 5*B*). Likewise, the AUC variance for different frequency bands of the wavelet-transformed data indicates a period of about 60 ms in the high-gamma frequency range (> 100 Hz) and another, rather long period in the alpha frequency range as candidate time-frequency windows of high sensitivity (Fig. 5*C*). Corresponding analyses were done for the wavelet terms *Re*{*W*} (Fig. 5*D*) and *Im*{*W*}, and phase angles (Fig. 5*E*). The selected epochs and frequencies were then used to trial-wise calculate the feature values across the array for various candidate features of both the EFP signal and its wavelet-transformed derivates, as illustrated in Fig. 4*B*.

### Classification performance for ROC-based feature selection

For each of the nine letter stimuli, Figure 6*A* shows the resultant high-gamma power (HGP) distribution over all the electrodes of M2 for each of 10 example trials per stimulus. Each electrode’s value represents power in a single trial, averaged over the most-informative time-frequency bins estimated by the ROC analysis. Although some electrodes were generally providing higher HGP than others, the specific pattern in response to each of the nine letters was quiet distinct. Training the SVM with these feature vectors yielded a mean single-trial classification performance of 87.2% based on the EFP, and 93.4% based on HGP, averaged over all stimulus conditions and monkeys. For HGP, this was a significant improvement (Wilcoxon signed rank, both *Z* = 4.35, *P* < 10^−4^, *R* = 0.081, *N* = 27) as compared to SVM performance based on the signals’ time course (cf. Fig. 3*E*, using the best electrode combinations per animal), and a corresponding statistical trend for the EFP (cf. Fig. 3*D, Z* = 1.73, *P* = 0.084). Additionally, ROC-based selection of specific time and time-frequency windows for extracting the values used in the distributed feature vectors outperformed whole trial-based value extraction for most of the response features we tested (Fig. 6*B*). HGP turned out to be very informative even when estimated by averaging all time bins of a trial, while all other features strongly benefitted from reducing the noise by discarding non-informative time bins. Thus, using the spatial distribution of signal features and ROC-based selection of time bins allowed to identify five response features with correct classification of more than 80% of single trials, averaged over all stimuli and arrays. Among these were three features derived from the wavelet-transformation (HGP, *Re*{*W*}, and *Im*{*W*}), and two features from the EFP (mean amplitude and max negativity).

**Fig. 6.**
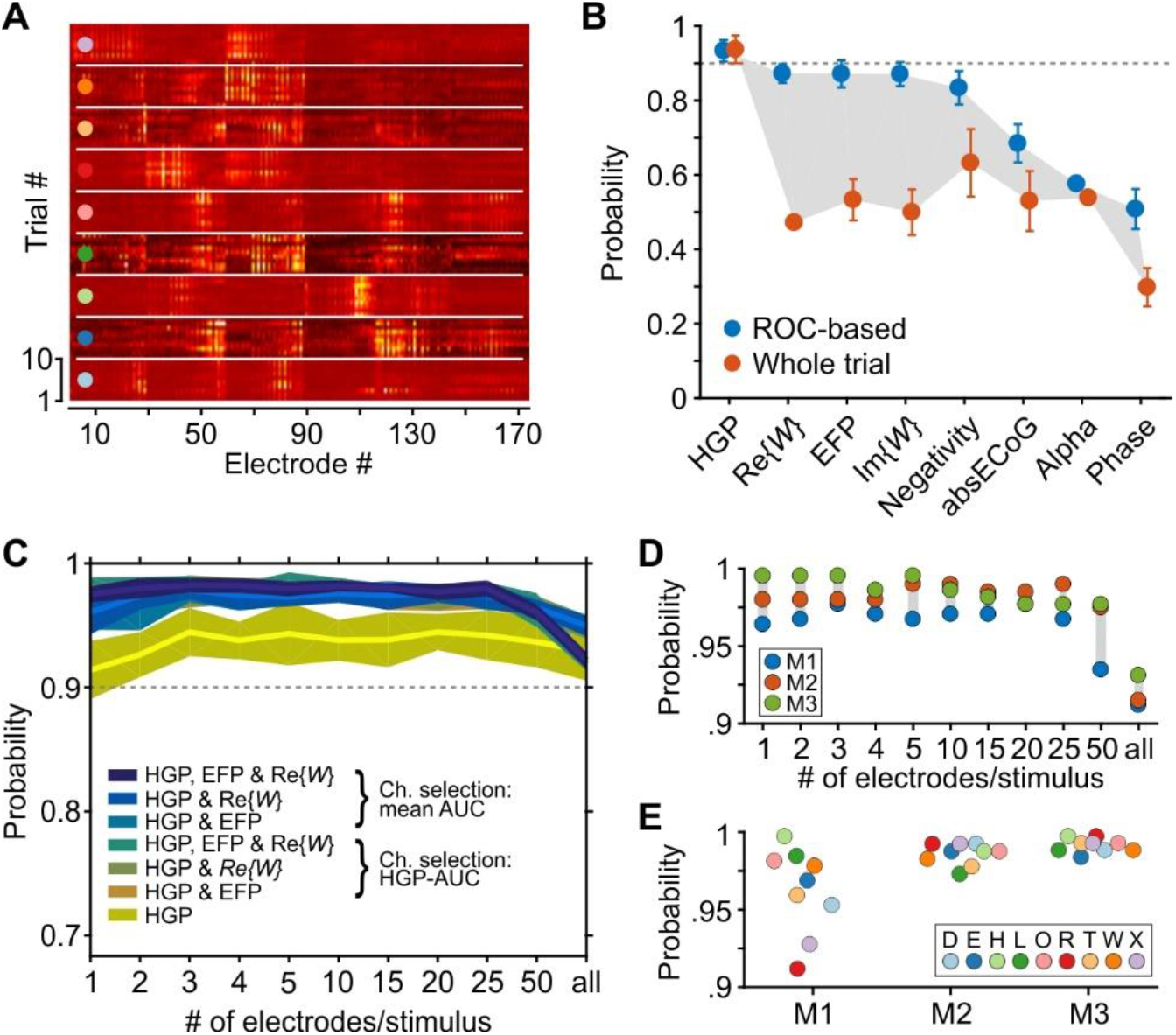
Identification of most-informative response features and electrodes. *A*: Example trials from M2, represented as the distributed activity of HGP over all electrodes of the array, sorted by stimulus condition. Only threshold-surpassing time-frequency bins as derived from preceding ROC analysis were considered for construction of the distributed activity vectors. *B*: Classification results for eight candidate features using the distributed activity vectors constructed from ROC-based selection of time (or time-frequency) windows, compared to classification using vectors constructed from all time bins of the trial (of the same frequency range, where applicable). *C*: Classification results for different combinations of the best response features as derived from *B*, for different numbers of electrodes used to construct the distributed activity vectors. Electrodes were selected either based on AUC variance for HGP, or based on the mean AUC variance of the combined features. *D*: Classification rates for the numerically best feature combination (HGP, *Re*{*W*}, and EFP amplitude, electrodes selected by mean AUC) as a function of number of electrodes used to construct the distributed activity vectors, individually for each animal. Each dot indicates median performance over the nine stimulus conditions. *E*: Classification rates for individual stimulus conditions based on the best selection of electrodes per animal. Note that in *D* and *E* the y-scale for probability is limited to the range of 0.9 to 1.

The response to each of the stimuli, however, was carried by only a small group of electrodes, while many electrodes did not respond, as expected due to the limited size of ERFs (Fig. 6*A*). Therefore, in analogy to selection of most-informative time bins, we next isolated most-informative electrodes for representation of distributed feature vectors, and discarded all others. To this end, we first chose the three features that allowed for best classification (HGP, *Re*{*W*}, and EFP amplitude). For classification, we started with the full set of electrodes and, per letter stimulus, either used a concatenated vector constructed from the mean HGP of each electrode, or alternatively, a concatenated vector using HGP plus either one or both of the other two features. Second, to reduce the number of electrodes, for each of the stimuli we stepwise excluded electrodes with the lowest AUC values and performed stimulus classification on vectors constructed from a decreasing number of electrodes with *N* = [50 25 20 15 10 5 4 3 2 1]. Third, to select most-informative electrodes we used two different procedures by either sorting electrodes by their HGP-AUC values only, or alternatively, by the mean AUC values across the two or three combined features. In summary, we tested different feature combinations on different sizes of electrode pools, and used different procedures to select the electrodes that remain in the pool. We thus ended up with 77 different data sets of feature values constructed from ROC-based selection of time and time-frequency windows, all of which were then tested for classifying single trials. The results are summarized in Figure 6*C*, showing the mean over the median performance per animal.

This analysis provided several insights: First, for nearly all of the data sets mean performance was clearly above our criterion level of 90% (Fig. 6*C*). Using only HGP as response feature, minimum median performance was 87.3% (M1 for electrode pool size of 1 per stimulus), and maximum median performance was 97.25% (M3 for electrode pool size of 5 per stimulus). Interestingly, adding information from at least one of the other two features (EFP and/or *Re*{*W*}) increased classification performance significantly (Kruskal-Wallis, *χ*^2^ =51.17, *P* < 10^−8^, *ω*^2^ = 0.2, *df* = 6). Post-hoc tests showed that HGP alone was worse than any other feature group (all *P* < 10^−3^), while none of the remaining groups was different from another (all *P* > 0.45). Second, there was no significant difference between the two procedures to select the most-informative electrodes (Wilcoxon signed rank, *Z* =1.404, *P* = 0.16, *R* = 0.007, *N* = 99). Both methods (selection either on HGP-AUC values or on mean AUC values of the concatenated features) worked equally well. Third, the number of electrodes used for classification made a significant difference (Kruskal-Wallis, *χ*^2^ = 45.23, *P* < 10^−5^, *ω*^2^ = 0.16, *df* = 10). Post-hoc tests revealed that using all electrodes to build the distributed activity vectors for each of the stimuli resulted in a significantly worse classification performance than using a limited number of more informative electrodes per stimulus (all *P* < 0.044 for < =25 electrodes). All other performance comparisons regarding the number of electrodes used for classification were not significantly different. Numerically, however, performance was poorest for the extreme groups (*N* < = 2 and *N* >= 50 electrodes per stimulus) and was higher and almost identical for pool sizes in-between (*N* = 3, 4, 5, 10, 15, 20 or 25 electrodes per stimulus, largest median performance difference: <0.35%). Figure 6*D* provides a corresponding example for the numerically best feature combination (concatenated vectors of HGP, *Re*{*W*} and EFP amplitude per electrode, and channel selection via HGP-AUC). Fourth, the high classification performance was true for each of the nine stimulus conditions in each of the three animals. When selecting the numerically best number of electrodes per animal, each individual stimulus condition was correctly classified in more than 90% of the trials. For 25 out of the 27 stimuli, classification reached the 95% performance level, and for 13 stimuli it even reached the 99% performance level (Fig. 6*E*). This was a highly significant performance increase as compared to the concatenated time course of the best-performing electrodes (cf. Fig. 3*E*, Best1paired condition) (Wilcoxon signed rank, *Z* = 4.54, *P* < 10^−5^, *R* = 0.084, *N* = 27). Thus the strong performance difference for the arrays of the three monkeys as seen when using the entire time course of the signal (cf. Fig. 3*D, E*) disappeared almost completely, and classification performance for the three arrays was statistically not different anymore (Kruskal-Wallis, *χ*^2^ = 3.75, *P* = 0.15, *df* = 2).

In summary, limiting the data to the most informative time (time-frequency) windows and electrodes by, in our case, simple ROC analysis, and combination of two or three response features allowed for superb classification rates of the EFP signal, proving that extremely short data fragments of visually driven, single-trial, epidurally recorded potentials convey highly localized information and represent the underlying functional architecture with high selectivity.

### Learning-free decoding and implications for future development of epidural arrays

Because selectivity and reliability of signal features are of outstanding importance for any scientific, technical, and clinical application using epi- or subdurally implanted arrays, we finally investigated the question to what extent design factors of the arrays influence decoding performance. For visually evoked signals, decoding performance will be directly related to a) sufficient activation of ERFs by the stimulus, b) robustness against small spatial displacements of the stimulus relative to the center of gaze, and c) robustness against signal variations. All of these factors place constraints for the electrode density of the array. This is illustrated by Figure 7*A* showing ERFs with a peak sensitivity within one degree of visual angle of the corresponding stimulus center for two example stimuli (data taken from M3). For stimulus R, the visual space around the letter was covered by a total of eight electrodes. This is expected to increase the probability to significantly activate one of these electrodes, regardless of the exact coordinates of the stimulus. Moreover, small displacements of the stimulus (e.g. caused by small eye movements) would not shift the stimulus out of the area covered by the electrodes but towards a neighboring ERF center (marked by crosses), and the denser ERF coverage provides some redundancy to compensate for electrodes with weaker signal-to-noise ratio. In contrast, stimulus D was covered by ERFs of only two electrodes, such that its neuronal representation is supposed to more strongly depend on the exact stimulus coordinates, and is likely more vulnerable against displacements of the stimulus and signal fluctuations.

**Fig. 7.**
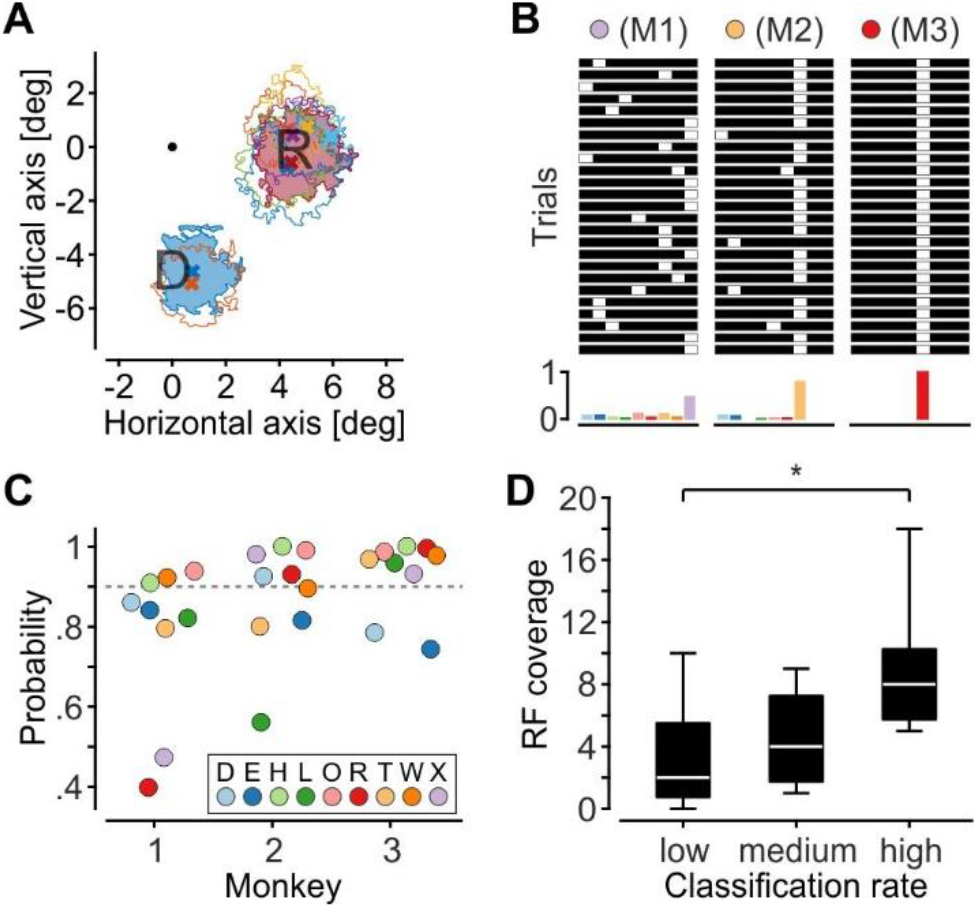
Relation between classification performance and ERF coverage. *A*: Receptive fields covering the location of two example stimuli, with ERF centers located within 1 deg radius of the letter center. Crosses indicate ERF centers. Filled areas indicate ERFs used for classification. Colors are randomly drawn and do not relate to stimulus conditions. *B*: Results of a binary classifier for 25 randomly drawn trials for three example stimuli (top panels). Each bar consists of nine elements, each representing the mean HGP of the best electrode per stimulus. The classifier assigns the value 1 to the element with the highest feature value (white square), and the value 0 to all other elements. Histograms (bottom panels) show the classification result for all trials of the stimuli. Color scheme is indicated in *C. C*: Single-trial classification results for each of the 3 * 9 stimulus conditions using the binary classifier. *D*: ERF coverage at stimulus locations as function of classification rates (lower, middle, and upper third of ranks). Boxes indicate 25^th^ and 75^th^ quartile, whiskers indicate full range of data, white bars indicate medians, and star indicates significant difference at the α = 5% level.

As shown in the preceding chapters, combining advanced machine learning algorithms and careful selection of informative data can compensate for some ambiguity in the neuronal representation of the stimuli. In fact, even the absence of a visual response may serve as a unique pattern to allow reliable classification of a stimulus. For lasting functionality and reliability, however, good coverage of the visual field and high density of ERFs is likely to be most critical.

This is illustrated by a re-classification of the 3 * 9 stimulus conditions with a simple binary classifier. For each array, the classifier was fed with a 9-element vector, where each element represented the single-trial HGP response averaged over the previously estimated most-informative time-frequency bins of the best electrode per letter stimulus. The element with the highest activation was set to one, while all others were set to zero (Fig. 7*B*), i.e. the decoding was based on a simple max function and independent of training a classifier. As expected, the overall classification performance of this extremely simple procedure significantly decreased as compared to the previous SVM-based classification (Wilcoxon signed rank, *Z* = 4.29, *P* < 10^−5^, *R* = 0.58, *N* = 27), yet single trials for 15 out of the 27 stimulus conditions were still classified with 90% or more correct, and two conditions reached the 100% performance level (Fig. 7*C*). When dividing all stimulus conditions into three equally large fractions associated with low, medium, and high classification performance, it was found that classification performance significantly (Kruskal-Wallis, *χ*^2^ = 8.01, *P* = 0.018, *ω*^2^ = 0.24, *df* = 2) increased as a function of ERF coverage (Fig. 7*D*). Because only one electrode per stimulus was used for decoding, this result is likely due to the fact that the array provides a raster of slightly shifted ERFs in regions of high electrode coverage, and consequently, a higher likelihood of good center-to-center alignment between stimulus and ERF. The results of the binary classifier show that under such conditions a very simple decoding strategy on an extremely small amount of data allow for close-to, or literally perfect, read-out of single-trial EFPs, proving that EFPs convey cortical signals with high spatial resolution and precisely reflect the underlying functional architecture of the cortex.

## Discussion

Recording of epidural field potentials, sometimes also termed □1EEG or epidural ECoG, has become an important technique for analysis of neuronal data at the mesoscopic level and offers promising options for future research on both basic and clinical neuroscience (Engel et al., 2005; Slutzky and Flint, 2017). Because electrodes are placed on top of the dura, EFP recordings are less invasive than subdural ECoG and intracortical recordings. This has the general advantage of a reduced risk of infection both during and after surgery (Van Gompel et al., 2008), and circumvention of other possible complications associated with opening the dura. On the other hand, the barrier of the dura may has an effect on spatial selectivity and signal quality, particularly when using electrodes with small diameter (Bundy et al., 2014). Both factors potentially constrain the usability of epidural signals for applications and research relying on clearly localized and/or single-trial information. We here show that despite these apparent drawbacks single-trial EFPs from three different arrays were classified in all of 27 stimulus conditions with a performance above 90%. For single trials from 25 out of the 27 conditions the performance level was more than 95%, and for 13 conditions it was larger than 99% of all trials. Hence, epidurally recorded EFP signals constitute a highly specific, local signal. Because we recorded in an area with high spatial resolution, the reliable classification of single trials suggests that the EFP is integrated over a clearly limited volume of cortical tissue, for otherwise spatial information would get blurred and errors in assigning the spatial condition would significantly rise. Importantly, reliable classification was not primarily dependent on advanced machine learning algorithms but could be achieved even with a binary classifier applying a simple max function. The latter result correlated with the electrode coverage of the cortical surface, indicating that a proper stimulus-ERF alignment is critical for the information content of the EFP.

### Methodological and technical considerations

Although we dedicated a good extent of the work to increase the classifier’s overall performance as a proxy for both the selectivity and reliability of the EFP signal, it is important to note that it was not the goal of the study to find, or suggest, the optimal procedure. Particularly for somatosensory cortex, much emphasis has been put on the optimization of classifiers and procedures, selection of signals, and choice of features, mostly using subdural or intracortical signals (Krusienski et al., 2011; Tankus et al., 2014; Slutzky and Flint, 2017). Yet, while some studies showed that also the EFP conveys sufficient information to be candidate as a source for BCI (Flint et al., 2012), the specific procedures that were applied, the experimental and clinical conditions, or the area delivering the neural signals usually not allowed to assess the functional specificity of the EFP. Epidural placement of electrodes introduces some additional variability, due to differences in dura thickness and volume of subdural space, slow changes in tissue impedance, and other factors. In the present study, this is exemplified by significant differences of both the size and SBR of ERFs between the arrays of the three monkeys, and across individual arrays. Together with differences in ERF coverage at different stimulus positions and strong trial-wise signal fluctuations, this variability predicts different probabilities for decoding stimulus information at the single-trial level, which constrains the use of EFPs. The goal of the present study, therefore, was to investigate the reliability by which these signals can be decoded. We showed that, whereas a difference between the arrays of the three monkeys and between individual stimuli was highly significant when decoding was based on the time course of the signal, an ROC-based approach for selecting the most-informative time bins and electrodes allowed to overcome these limitations and to classify single trials for all of 27 conditions with 90% correctness, at the least. Because this analysis was performed in an area with highly resolved retinotopic representation, our results prove that EFP recordings provide a strongly localized signal that can be directly related to the functional architecture of the underlying tissue, even when information is limited to single trials. This has important implications not only for the analysis of EFPs but also for the interpretability of subdurally recorded ECoG data derived from cortical areas having a less clear, or more dynamic, functional organization than V1.

Even though optimization of the classification procedure was not the primary focus of the study, our results allow, nevertheless, for some general methodological and technical conclusions. First, as reported before (Rotermund et al., 2009; Gunduz et al., 2012; Kapeller et al., 2018; Martin et al., 2019), visual gamma activity was found to be an excellent feature for distinguishing between different stimulus conditions. Yet, combining gamma with one or two other highly informative response features allowed for significant improvement of classification performance. Similar conclusions were drawn by other authors using combinations of different response features (Wei et al., 2007; Zhang et al., 2013; Nourski et al., 2015; Miller et al., 2016; Li et al., 2017). This finding is likely indicating that, if they are sufficiently independent, ambiguity in the information of one feature might be compensated by disambiguity in another. Second, excluding weakly or non-informative data by using only the most-informative time (time-frequency) windows and electrodes not only reduced the computational load by orders of magnitude but significantly improved classification performance. In line with our findings in visual cortex, studies in somatosensory cortex reported improved, or equally well, decoding performance by reducing the number of electrodes to some lower limit (Wei et al., 2010; Pan et al., 2018), and by referring to only a few time bins (Kaiju et al., 2017). We performed selection of most-informative electrodes and time-frequency bins by ROC, but other procedures for distinguishing between distributions of data from different conditions may work equally well. Third, classification performance correlated with ERF coverage of the cortical surface. This is likely due to the fact that a larger number of candidate electrodes significantly increases the likelihood to identify one or a few highly selective channels, whereas low coverage of the cortical surface induces a strong dependence on the signal quality of a limited number of available channels. In line with this, studies in somatosensory cortex reported high classification rates when using high-density ECoG arrays to decode hand and finger gestures based on information from a single channel per condition (Kaiju et al., 2017).

### Functional specificity of epidural field potentials

Many of the studies recording signals from top of the brain have been performed using subdural arrays with large electrodes (Crone et al., 2006; Slutzky and Flint, 2017). Epidural arrays, however, offer a less traumatic access to intracranial brain signals, and commercially available arrays have recently been used for e.g. diagnosis of the cognitive state in locked-in syndrome (Martens et al., 2014; Bensch et al., 2014) and rehabilitation of patients after stroke (Gharabaghi et al., 2014a; 2014b). In the motor system, recent studies in non-human primates showed that EFPs allow well decoding of hand and finger gestures (Choi et al., 2018). In the visual domain, monkey EFPs were used to study gamma band responses (Taylor et al., 2005; Rotermund et al., 2009; Grothe et al., 2012), to predict allocation of attention between two close-by locations (Rotermund et al., 2013), and to predict saccade directions (Lee et al., 2017). Yet, probably because of the large number of neurons contributing to the EFP/ECoG signal (Miller et al., 2009), the additional attenuation of the signal by the dura makes epidural recordings a still rare case, despite its obvious clinical advantages. Two recent studies in monkey somatosensory cortex indeed indicated weaker decoding performance for epidural as compared to subdural signals (Bundy et al., 2014; Farrokhi and Erfanian, 2018). On the other hand, a rodent study on forelimb movements concluded that EFPs allow for the same decoding performance than intracortical LFPs (Slutzky et al., 2011). In fact, in-vivo measurements of signal attenuation by human dura indicated no detrimental effects on signal feature detection (Torres Valderrama et al., 2010). A more significant factor determining the specificity of EFP/ECoG signals, therefore, might be given by the size and density of the recording electrodes. Only few studies have compared signal quality and decoding performance for the different types of subdural and intracortical arrays used in clinical and basic neuroscience research (Kellis et al., 2016; Flint et al., 2017; Wang et al., 2017), yet it appears that many of the results indicating high signal specificity were obtained with high-density arrays and rather small electrodes (Kaiju et al., 2017; Branco et al., 2017; Hu et al., 2018; Ramsey et al., 2018). The findings of the present study that EFPs possess high spatial selectivity is clearly in line with this: With the small electrodes of our array we estimated mean ERF sizes of about 2.7 and 2.8 deg diameter (re-calculated from ERF areas assuming circular ERFs) in monkeys M1 and M3, respectively, and 4.0 deg in M2, which is around a factor of 1.6 to 2.3 of the receptive field size of intracortical LFPs, measured with the same mapping paradigm (Drebitz et al., 2019). A recent study using large, subdural electrodes reported about the same factor when comparing ECoG to LFP ERFs (Dubey and Ray, 2019). Two other studies using subdural electrodes estimated V1-RFs of slightly smaller (Yoshor et al., 2006) and of about the same size we here report for epidural electrodes (Takaura et al., 2016). Thus, regarding selectivity of the signal, sub- or epidural placement of electrodes may not be the most limiting factor. In fact, because we achieved very good single-trial classification of many stimulus locations even without training advanced classifiers, using multielectrode arrays with small electrode diameter and high electrode density seems critical for improving the reliability by which single-trial information can be read out from both sub- and epidural recordings. This is also supported by a recent study in ferrets using high-density arrays with electrodes of even smaller diameter than ours (Bockhorst et al., 2018). The authors compared epicortical recordings with intracortical laminar probe recordings and found well-preserved receptive field properties in epicortical recordings, probably due to the fact that epicortical signals reflect the highly synchronized cortical activity. Although epidural placement of electrodes increases the distance between electrodes and cortical sources, and likely reduces the correlations between epi- and intracortical signals, small electrodes will compensate for the larger distance by integrating over a smaller volume of tissue, thus providing selective estimates of cortical activity.

Taken together, despite strong trial-by-trial fluctuations, and signal attenuation by the dura, epidurally recorded signals convey a very high degree of spatial selectivity that is reliably available at the single-trial level. This offers new and exciting options for both clinical and basic neuroscience, and for using EFPs as the source of information in brain-computer interfacing.

## Author contributions

Conceptualization: B.F. and D.W.; Methodology: B.F., A.K.K., A.S., W.L., and D.W.; Investigation: B.F.; Data Analysis: B.F. and D.W.; Figures: B.F. and D.W.; Writing – Original Draft: B.F. and D.W.; Writing – Review & Editing: B.F., A.K.K., A.S., W.L., and D.W.

## Acknowledgments

The work was funded by grants from DFG (WE 5469/3-1), Tönjes-Vagt Foundation (Project XXXI), University of Bremen (ZF grant & Creative Unit I-SEE), and a scholarship from the German Academic Scholarship Foundation. The authors declare no competing financial interests. The authors thank Udo Ernst for helpful comments on an earlier draft and Peter Bujotzek, Katja Taylor, Serge Strokov, Ramazani Hakizimani, and Katrin Thoß for technical support.

